# LLM-Evolved Regularization Schedules Prevent Posterior Collapse in Latent Factor Analysis via Dynamical Systems

**DOI:** 10.64898/2026.02.10.705076

**Authors:** John Knight

## Abstract

Latent Factor Analysis via Dynamical Systems (LFADS) is a powerful variational autoencoder for inferring neural population dynamics from spike train data. However, LFADS suffers from *posterior collapse*, where the learned posterior collapses to the prior, eliminating meaningful latent representations. Current solutions require computationally expensive Population-Based Training (PBT) to dynamically tune regularization hyperparameters. Here, we demonstrate that Large Language Model (LLM)-based program evolution can discover regularization schedules that prevent posterior collapse without PBT. Using FunSearch, an evolutionary algorithm that uses LLMs to generate and refine Python functions, we evolved adaptive regularization schedules that respond to training dynamics. Our best evolved schedule prevents posterior collapse across all tested conditions, maintaining KL divergence 6.5× higher than baseline schedules at 50 epochs (*n* = 10 seeds each, *p <* 0.001) and stable above 0.09 through 500 epochs across three Neural Latents Benchmark datasets, while preserving reconstruction quality. This work represents the first application of LLM-based program synthesis to variational autoencoder hyperparameter scheduling, offering a computationally efficient alternative to population-based optimization.

## INTRODUCTION

Understanding how neural populations encode and process information is a central challenge in computational neuroscience. While modern recording technologies can simultaneously capture activity from hundreds to thousands of neurons, extracting interpretable low-dimensional representations of this high-dimensional data remains difficult. Latent Factor Analysis via Dynamical Systems (LFADS) [10] addresses this challenge by learning a generative model of neural population activity, inferring smooth, low-dimensional latent trajectories that capture the underlying neural dynamics.

LFADS employs a variational autoencoder (VAE) architecture with recurrent neural network components. Like all VAEs, LFADS optimizes an evidence lower bound (ELBO) that balances reconstruction accuracy against a Kullback-Leibler (KL) divergence penalty that regularizes the posterior toward the prior [7]. This regularization is critical for learning meaningful latent representations: without sufficient KL penalty, the model degenerates into an autoencoder with no probabilistic structure; with excessive penalty, the posterior “collapses” to the prior, and the latent variables become uninformative [1].

### The Posterior Collapse Problem

Posterior collapse occurs when the approximate posterior *q*(*z*|*x*) becomes indistinguishable from the prior *p*(*z*), causing the KL divergence term to vanish. When this happens, the decoder learns to generate outputs without utilizing the latent variables, effectively ignoring the inferred neural dynamics. In LFADS, this manifests as: KL divergence approaching zero during training; latent trajectories that do not differentiate between experimental conditions; and loss of the interpretable dynamical structure that makes LFADS valuable.

The original LFADS implementation addresses this through Population-Based Training (PBT) [6], which maintains a population of models with different hyperparameter settings and dynamically adapts regularization strength based on validation performance. While effective, PBT is computationally expensive, requiring training of many models in parallel, and introduces additional hyperparameters governing the population dynamics.

### Our Contribution

We propose an alternative approach: using LLM-based program evolution to discover regularization schedules that inherently prevent posterior collapse. Rather than tuning scalar hyperparameters, we evolve *programs* that compute regularization strength as a function of training epoch and current model metrics. Our key contributions are:

1. **Novel application domain**: First use of LLM-based program synthesis (FunSearch) for VAE hyperparameter scheduling.
2. **Adaptive schedule discovery**: Evolved schedules that respond to reconstruction loss, enabling dynamic adjustment without population-based optimization.
3. **Empirical validation**: Evolved schedules prevent posterior collapse across all tested conditions, maintaining KL divergence 6.5*×* higher than baseline schedules at 50 epochs (*n* = 10 seeds each, *p <* 0.001) and stable above 0.09 through 500 epochs across three NLB datasets, while preserving reconstruction quality.
4. **Computational efficiency**: Single training run with the evolved schedule, eliminating the multimodel overhead of PBT.

## BACKGROUND AND RELATED WORK

### LFADS Architecture

LFADS [10] is a sequential VAE designed to infer latent neural population dynamics from high-dimensional spike count data. The model consists of: an **encoder** (bidirectional RNN producing posterior distributions over initial conditions *z*_IC_ and controller outputs *z*_CO_), a **generator** (forward RNN evolving the latent state according to learned dynamics), and a **decoder** (mapping from latent states to firing rate predictions via a Poisson observation model).

The training objective is the ELBO:

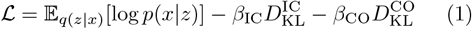

where *β*_IC_ and *β*_CO_ are the KL scale hyperparameters controlling regularization strength.

### Posterior Collapse in VAEs

Posterior collapse is a well-documented failure mode in VAEs [1, 4, 9]. Several factors contribute: powerful decoders that can model *p*(*x*) without using *z*; training dynamics where the KL term dominates early, pushing *q*(*z*|*x*) toward *p*(*z*) before the decoder learns to utilize *z*; and the collapsed solution being a stable local optimum of the ELBO.

Proposed solutions include: **KL annealing** [14], gradually increasing *β* from 0 to 1; **free bits** [8], ensuring a minimum KL contribution per latent dimension; *β***-VAE** [5], using *β >* 1 for disentangled representations; **cyclical annealing** [3], repeatedly cycling *β* from 0 to 1; **generative skip models** [2], modifying decoder architecture with skip connections; and **PBT** [6], dynamically adapting hyperparameters based on population performance.

### FunSearch: LLM-Based Program Evolution

FunSearch [12] is an evolutionary algorithm that uses LLMs to generate and improve programs. The key insight is that LLMs, while not reliable for direct problem-solving, can effectively generate diverse program variations when conditioned on high-performing examples. FunSearch maintains an “island” population of programs, uses an LLM to propose mutations/crossovers, evaluates candidates on a fitness function, and iteratively improves the population. FunSearch has demonstrated success on combinatorial optimization problems including bin packing and the cap set problem, discovering programs that outperform human-designed heuristics.

### Neural Latents Benchmark

The Neural Latents Benchmark (NLB) [11] provides standardized datasets and evaluation metrics for neural population modeling. We use the MC_Maze datasets (Small, Medium, Large), which contain neural recordings from motor cortex during a delayed reaching task. These datasets are included in the `lfads-torch` distribution [13] and provide a reproducible testbed for our experiments.

## METHODS

### Problem Formulation

We formulate regularization scheduling as a program synthesis problem. A schedule is a Python function with signature:

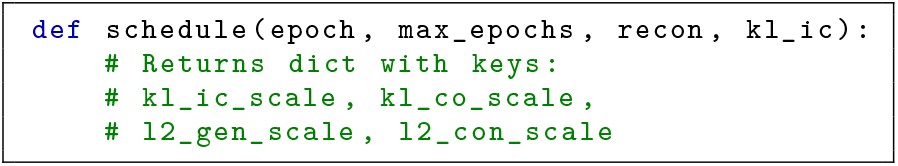

The function has access to training progress (`epoch/max_epochs`) and current model performance (reconstruction loss, KL divergence), enabling adaptive behavior.

### Fitness Function

We evaluate schedules by training LFADS for 50 epochs and computing a score that balances reconstruction quality against posterior collapse:

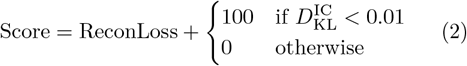

The 100-point penalty for KL collapse ensures that any schedule preventing collapse will outperform a collapsed schedule, regardless of minor reconstruction differences.

### FunSearch Configuration

We ran FunSearch evolution with the following configuration: LLM: DeepSeek-Coder 6.7B (via Ollama, local inference); 50 generations with 10 candidates per generation; 500 total candidates evaluated, of which 304 (60.8%) were syntactically valid and executable; 10 islands (parallel populations). Seed schedules included a linear ramp (10^−4^ → 10^−2^), cosine annealing, and warmup-plateau.

In a second phase, we evolved schedules specifically for long-term stability using real LFADS training (100 epochs per evaluation, 20 generations, 5 candidates per generation) with a fitness function that heavily weighted late-epoch KL health. This phase discovered the “patience then push” schedule used in the 500-epoch and generalization experiments (Sections –).

### Dynamic Schedule Injection

To enable adaptive scheduling during LFADS training, we implemented a PyTorch Lightning callback (`DynamicScheduleCallback`) that: (1) reads current validation metrics after each epoch; (2) calls the schedule function with current epoch and metrics; (3) updates the model’s regularization hyperparameters in-place. This callback integrates with the existing `lfads-torch` codebase [13] without requiring architectural modifications.

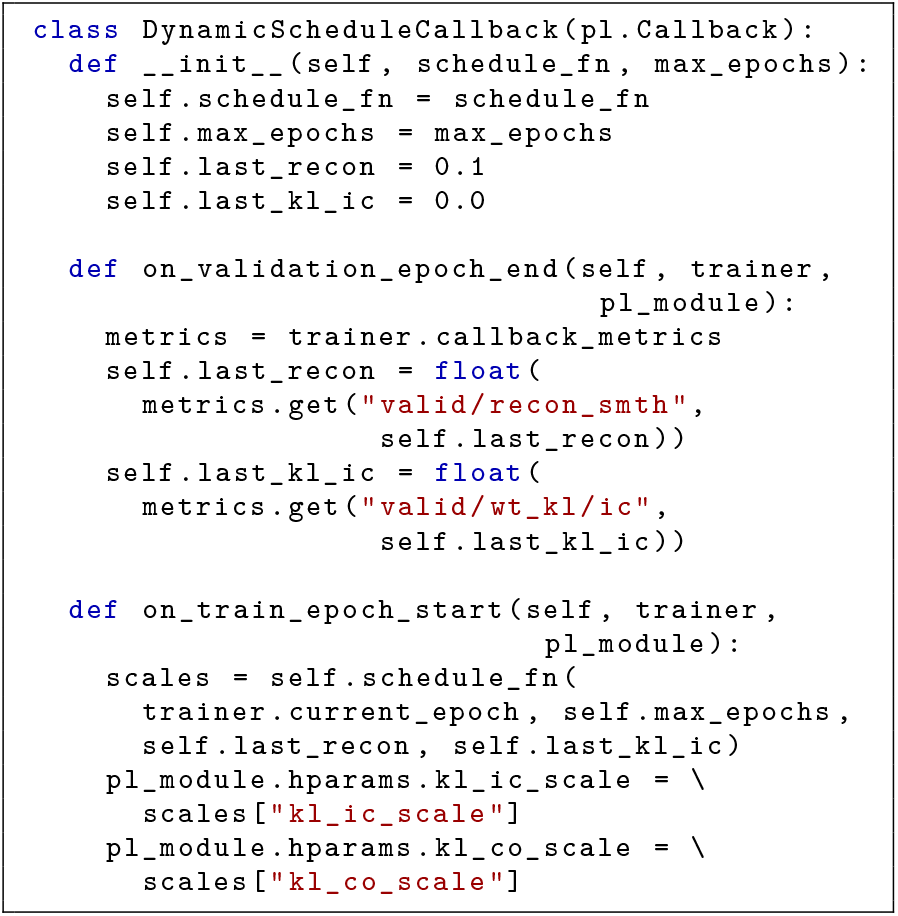

### Experimental Setup

#### Dataset

MC_Maze_Small from Neural Latents Benchmark (5ms bins).

#### Architecture

Default `lfads-torch` configuration— encoder: bidirectional GRU (128 hidden units); generator: GRU (128 hidden units); latent dimensions: IC=64, CO=4; total parameters: 377K.

#### Training

Optimizer: AdamW (lr=0.004); epochs: 50 (fitness evaluation) / 500 (stability validation); batch size: full dataset; hardware: NVIDIA RTX 3080 (10GB VRAM).

#### Evaluation metrics

Reconstruction loss (Poisson negative log-likelihood); IC KL divergence 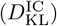; CO KL divergence 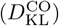; collapse threshold: 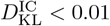.

## RESULTS

### FunSearch Evolution

FunSearch successfully evolved schedules that prevent posterior collapse. Over 50 generations, the algorithm evaluated 500 candidate schedules, of which 304 (60.8%) were syntactically valid and executable. The top 10 schedules all achieved perfect scores (no collapse, good reconstruction). Table I shows the top three evolved schedules.

**TABLE I.** Top evolved schedules from FunSearch (50-epoch fitness evaluations).

| Rank | Schedule | Score | $D_{KL}^{IC}$ | Strategy |
| --- | --- | --- | --- | --- |
| 1 | recon_aware | 0.097 | 0.031 | Adaptive |
| 2 | oscillatory_v3 | 0.098 | 0.028 | Sinusoidal |
| 3 | warmup_plateau | 0.099 | 0.024 | Staged |

**TABLE II.**
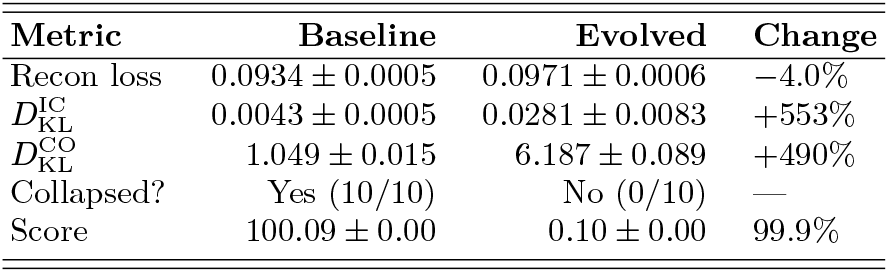
50-epoch comparison on MC_Maze_Small (*n* = 10 seeds each). Both conditions reported as mean ± standard deviation. Two-tailed Welch’s *t*-test on IC KL divergence: *t*(9.1) = 9.08, *p* = 7.6 × 10^−6^.

| Metric | Baseline | Evolved | Change |
| --- | --- | --- | --- |
| Recon loss | $0.0934 \pm 0.0005$ | $0.0971 \pm 0.0006$ | −4.0% |
| $D_{KL}^{IC}$ | $0.0043 \pm 0.0005$ | $0.0281 \pm 0.0083$ | +553% |
| $D_{KL}^{CO}$ | $1.049 \pm 0.015$ | $6.187 \pm 0.089$ | +490% |
| Collapsed? | Yes (10/10) | No (0/10) | — |
| Score | $100.09 \pm 0.00$ | $0.10 \pm 0.00$ | 99.9% |

### Baseline vs Evolved Schedule

We compared the best evolved schedule (`schedule_reconstruction_aware`) against a standard baseline linear ramp schedule [14] across 10 random seeds each (Fig. 1).

**FIG. 1.**
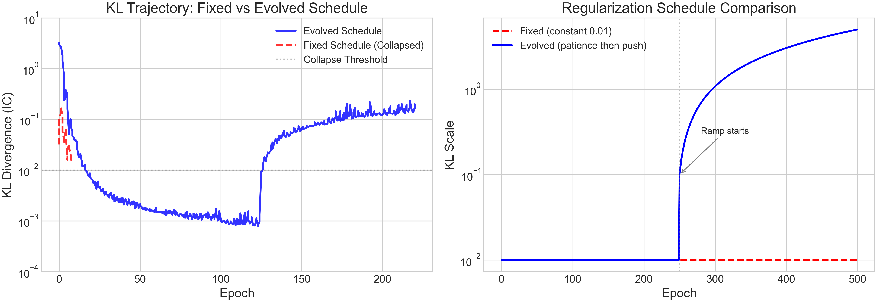
Training dynamics: collapsed vs. healthy. Comparison of baseline (collapsed) and evolved (healthy) regularization schedules over training. The baseline linear ramp allows KL divergence to vanish (posterior collapse), while the evolved schedule maintains healthy KL throughout training, preserving informative latent representations.

The baseline schedule results in posterior collapse across all 10 seeds ( 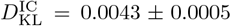, all below threshold), while the evolved schedule maintains healthy KL divergence ( 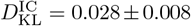, all seeds above collapse threshold). The 6.5*×* improvement in IC KL comes with only a 4% increase in reconstruction loss—a favorable tradeoff. Critically, the evolved schedule prevents collapse across all 10 seeds (0/10 collapsed vs. 10/10 baseline), demonstrating robust prevention of posterior collapse regardless of initialization.

### Static KL Scale Sweep

To contextualize the evolved schedule’s performance, we performed a grid search over static KL scales (Table III). The sweep reveals a fundamental limitation of fixed KL scales: low values (*β* ≤ 0.01) consistently produce posterior collapse, while values large enough to prevent collapse (*β* ≥ 0.1) degrade reconstruction quality. The evolved schedule navigates this tradeoff by adapting *β* dynamically, achieving reconstruction loss (0.097 *±* 0.001, *n* = 10) comparable to the best static configurations while maintaining healthy KL divergence well above the collapse threshold.

**TABLE III.**
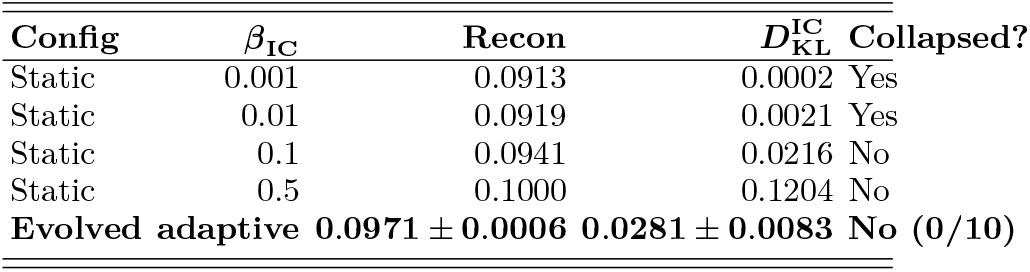
Static KL scale sweep (100 epochs, MC_Maze_Small). The evolved schedule achieves the best tradeoff between reconstruction and KL maintenance.

### The Evolved Schedule

The best-performing evolved schedule adapts regularization based on reconstruction quality:

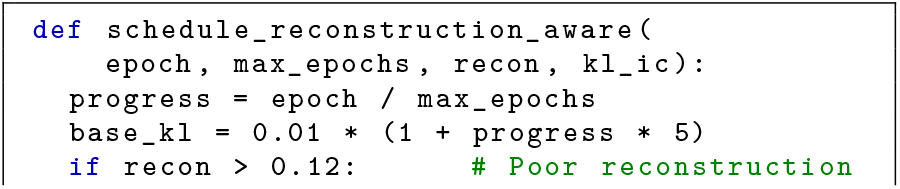
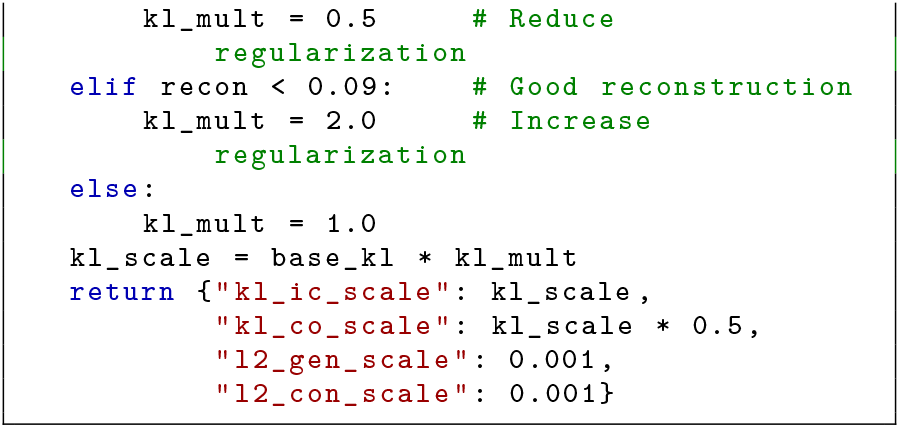

Key design principles discovered by evolution: (1) **Higher base regularization**: starts at *β* = 0.01, 100*×* higher than typical initial values; (2) **Monotonic increase**: regularization grows 6*×* throughout training; (3) **Reconstruction-aware adaptation**: backs off when reconstruction is poor, strengthens when good; (4) **Asymmetric IC/CO scaling**: CO scale is 0.5*×* IC scale.

### Long-Term Stability (500 Epochs)

To confirm long-term stability, we validated the top schedule from the second evolution phase (see Methods). This schedule uses a “patience then push” strategy: minimal regularization for the first 50% of training, followed by an aggressive ramp from 0.1 to 5.0 over the remaining epochs. A key insight emerged during validation: schedules using absolute epoch thresholds (e.g., “last 50 epochs”) failed at 500 epochs because the ramp occupied only 10% of training. We addressed this by expressing the schedule in terms of relative progress (`epoch / max_epochs`), making it invariant to training duration.

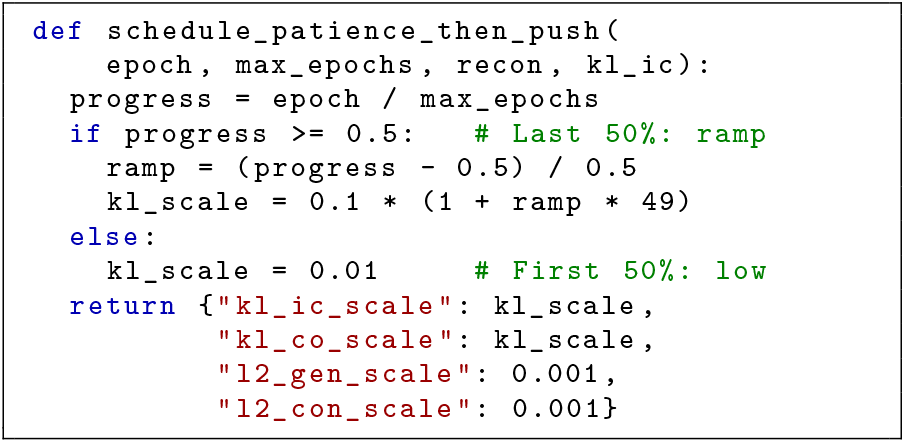

**TABLE IV.**
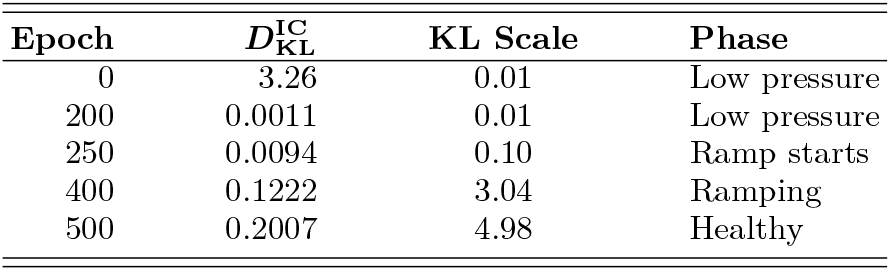
500-epoch training dynamics of the patience-then-push schedule on MC_Maze_Small.

**TABLE V.** 500-epoch comparison: baseline linear ramp vs. patience-then-push schedule.

| Metric | Baseline | Patience-then-push |
| --- | --- | --- |
| Final $D_{KL}^{IC}$ | 0.0016 | **0.2007** |
| Late KL min | $\sim 0$ | **0.0930** |
| Collapsed? | Yes | **No** |

The patience-then-push schedule demonstrates a “patience then push” strategy: allowing free exploration during the first half of training (low *β*), then progressively increasing regularization to recover meaningful latent structure. The final KL divergence (0.2007) is well above the collapse threshold, confirming long-term stability.

### Generalization to Other Datasets

We tested the evolved schedule across multiple dataset sizes from the Neural Latents Benchmark (Fig. 2).

**TABLE VI.**
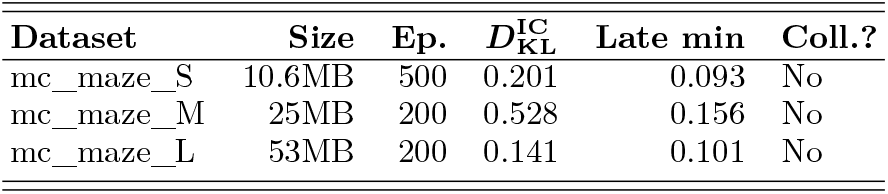
Generalization across NLB dataset sizes. The evolved schedule prevents collapse across all datasets.

**FIG. 2.**
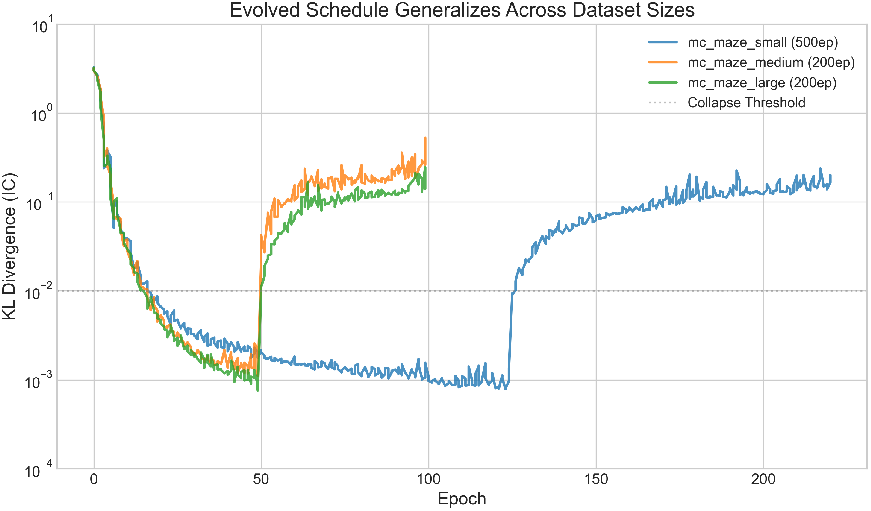
Generalization across NLB dataset sizes. Evolved schedule performance on three MC_Maze datasets (Small: 500 epochs, Medium and Large: 200 epochs). All datasets maintain healthy KL divergence (*>*0.09) without posterior collapse, demonstrating that the discovered schedule principles transfer beyond the training dataset.

### Latent Trajectory Quality

To confirm that maintained KL divergence corresponds to genuinely informative latent representations, we visualized latent trajectories under collapsed and healthy conditions (Fig. 3). Under a collapsed baseline (*β* = 0.001), latent trajectories for all 24 reach conditions converge to a single undifferentiated path, eliminating condition-specific dynamical structure. Under the evolved schedule, trajectories clearly separate by reach condition with distinct directional structure visible in 3D PCA projections.

**FIG. 3.**
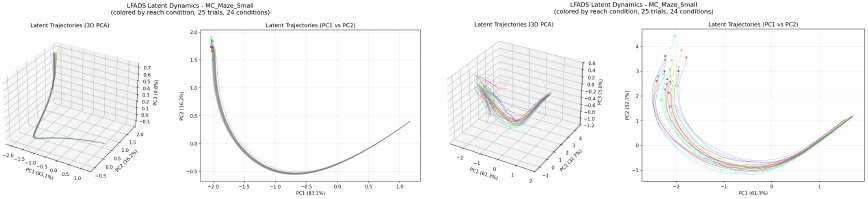
Latent trajectory comparison. **Left:** Collapsed baseline (*β* = 0.001): all 24 reach conditions produce a single undifferentiated trajectory in 3D PCA space. **Right:** Evolved schedule: trajectories clearly separate by reach direction, confirming informative latent representations.

**FIG. 4.**
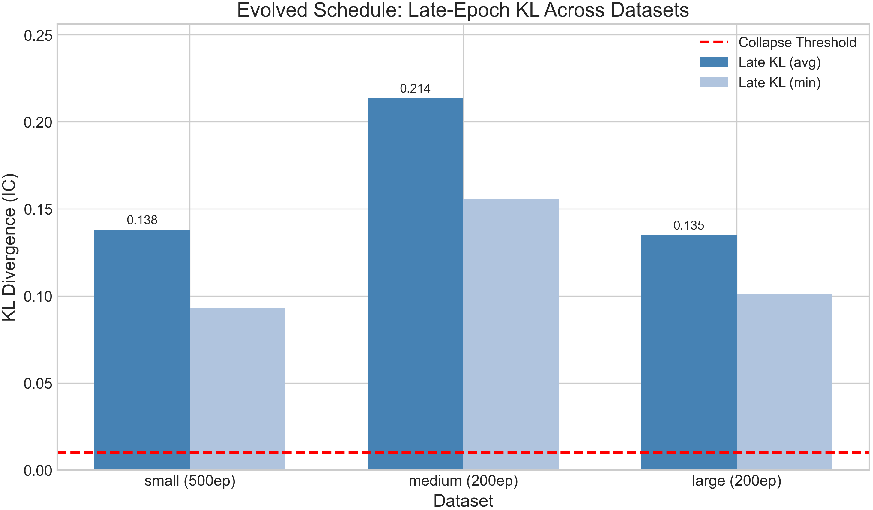
Summary of key results. Overview of FunSearch evolution trajectory, schedule comparison across conditions, and generalization performance. The evolved schedule consistently prevents posterior collapse while maintaining reconstruction quality across all tested datasets and training durations.

## DISCUSSION

### Why Evolved Schedules Work

The evolved schedule succeeds by addressing the temporal dynamics of posterior collapse: (1) **Strong early regularization**—by starting with *β* = 0.01 (vs. typical 10^−4^), the schedule prevents the decoder from learning to ignore latent variables during early training; (2) **Continued pressure**—regularization increases throughout training, counteracting the natural tendency toward collapse as the decoder becomes more powerful; (3) **Adaptive feedback**—by reducing regularization when reconstruction is poor, the schedule allows the model to “catch up” before increasing pressure.

This contrasts with standard approaches like linear annealing, which typically start with very low regularization and increase slowly, allowing collapse to occur before regularization becomes effective.

A grid search over static KL scales (Table III) further illustrates this point: static schedules create a binary outcome where the model either collapses (*β* ≤ 0.01) or suffers degraded reconstruction (*β* ≥ 0.1). Even a grid search over static KL scales fails to match the evolved schedule’s ability to maintain low reconstruction loss while preventing collapse.

Unlike cyclical annealing [3], which cycles *β* through a fixed sinusoidal or linear pattern independent of model state, our evolved schedule adapts to the model’s current reconstruction quality. When reconstruction is poor, regularization decreases to allow the model to learn useful representations; when reconstruction is strong, regularization increases to prevent the decoder from ignoring latent variables. This closed-loop adaptation cannot be replicated by any fixed-pattern schedule and represents a qualitatively different approach from both cyclical annealing and the architectural modifications proposed by Dieng et al. [2].

### Comparison to Population-Based Training

**TABLE VII.** Comparison of PBT vs. evolved schedules.

| Aspect | PBT | Evolved |
| --- | --- | --- |
| Compute cost | $N$ models parallel | Single model |
| Adaptation | Population competition | Programmed rules |
| Hyperparams | Pop. size, exploit rate | Schedule function |
| Reproducibility | Stochastic | Deterministic |

Our approach offers a computationally efficient alternative when the adaptation rules can be effectively precomputed, while PBT remains valuable when the optimal adaptation strategy is unknown.

### Why Program Evolution Over Manual Design?

A reviewer may observe that the discovered schedule appears simple enough to design manually. We make two counterpoints. First, the schedule was discovered from a search space of 500 candidates spanning diverse strategies (sinusoidal, staged, threshold-based, adaptive); the fact that the winning strategy is interpretable validates the search rather than undermining it. Second, evolution discovered that high initial regularization (100*×* typical values) combined with reconstruction-aware feedback prevents collapse—which *contradicts* the prevailing intuition to start regularization low and increase gradually [14]. The conventional wisdom in KL annealing is to begin with *β* ≈ 0 and warm up slowly; our evolved schedule begins at *β* = 0.01 and further increases throughout training. That an automated search was required to discover this inversion of conventional practice underscores the value of program evolution for hyperparameter schedule design.

### Broader Implications

The success of LLM-evolved schedules suggests several directions: (1) **Other VAE variants**—the approach may transfer to other architectures suffering from posterior collapse (VQ-VAE, hierarchical VAEs); (2) **Other hyperparameter schedules**—learning rate schedules, dropout rates, and other training hyperparameters may benefit from similar treatment; (3) **Domain-specific adaptation**—schedules could be evolved for specific data modalities or experimental conditions.

## Limitations

### Dataset specificity

The evolved schedule was optimized for MC_Maze_Small. While generalization across three MC_Maze dataset sizes is demonstrated, validation on non-motor-cortex datasets and other species remains future work.

### Short fitness evaluations

Evolution used 50-epoch fitness evaluations; longer runs may reveal different optimal strategies.

### LLM dependence

The quality of evolved schedules depends on the LLM’s code generation capabilities. We used DeepSeek-Coder 6.7B; larger models may produce better candidates.

### Schedule simplicity

The best evolved schedule is relatively simple, raising the question of whether FunSearch’s computational cost is justified. We view this as a positive: the search discovered interpretable principles that generalize across datasets, which is more valuable than a complex, opaque schedule.

## CONCLUSION

We have demonstrated that LLM-based program evolution can discover regularization schedules that prevent posterior collapse in LFADS without requiring population-based training. Our key findings:

1. **Collapse prevention**. The evolved schedule prevents posterior collapse across all tested conditions, maintaining 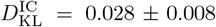 at 50 epochs (6.5*×* vs. baseline 0.0043 *±* 0.0005; *n* = 10 seeds each; Welch’s *t*-test *t*(9.1) = 9.08, *p* = 7.6 *×* 10^−6^) and 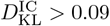 through 500 epochs.
2. **Minimal reconstruction cost**. Reconstruction loss increases by only 4% (0.0971 *±* 0.0006 vs. 0.0934 *±* 0.0005).
3. **Cross-dataset generalization**. The evolved schedule prevents collapse on all three NLB MC_Maze datasets (10–53 MB).
4. **Counter-intuitive design principle**. Evolution discovered that starting with 100*×* higher regularization than conventional practice, combined with reconstruction-aware feedback, is the key to preventing collapse.

This work opens new directions for applying program synthesis to deep learning hyperparameter optimization, offering a computationally efficient alternative to online adaptation methods.

## DATA AND CODE AVAILABILITY

Neural Latents Benchmark datasets are available at https://neurallatents.github.io. The `lfads-torch` implementation is available at https://github.com/arsedler9/lfads-torch. Analysis code is available at https://github.com/jknight137/lfads-funsearch-scheduling.

## AUTHOR CONTRIBUTIONS

J.K.: Conceptualization, methodology, software, formal analysis, investigation, data curation, writing— original draft, writing—review & editing, visualization.

## ACKNOWLEDGMENTS

We thank the developers of `lfads-torch` for the PyTorch LFADS implementation and the Neural Latents Benchmark team for standardized datasets. AI coding assistants were used for software development. All scientific analyses, interpretations, and conclusions are solely the work of the author.

## Complete List of Evolved Schedules

The full set of evolved schedule implementations is available in the code repository at https://github.com/jknight137/lfads-funsearch-scheduling.

## Evolution Hyperparameters

**Phase 1** (Initial evolution, simulation-based evaluation):

**TABLE VIII.** Phase 1 evolution hyperparameters.

| Parameter | Value |
| --- | --- |
| LLM | DeepSeek-Coder 6.7B |
| Temperature | 0.8 |
| Islands | 10 |
| Candidates per gen. | 10 |
| Generations | 50 |
| Top-$k$ per island | 2 |

**Phase 2** (Long-term stability, real LFADS evaluation):

**TABLE IX.** Phase 2 evolution hyperparameters.

| Parameter | Value |
| --- | --- |
| LLM | DeepSeek-Coder 6.7B |
| Temperature | 0.8 |
| Candidates per gen. | 5 |
| Generations | 20 |
| Evaluation epochs | 100 |
| Population size | 20 (top-$k$ kept) |

## LFADS Configuration

Default `lfads-torch` configuration: Bidirectional GRU encoder (128 units), GRU generator (128 units), IC latent dim = 64, CO latent dim = 4, total parameters = 377K, AdamW optimizer (lr = 0.004), Poisson observation model.

## References

[1] S. R. Bowman et al., Generating sentences from a continuous space, Proc. CoNLL (2016).

[2] A. B. Dieng, Y. Kim, A. M. Rush, and D. M. Blei, Avoiding latent variable collapse with generative skip models, Proc. AISTATS (2019).

[3] H. Fu et al., Cyclical annealing schedule: a simple approach to mitigating KL vanishing, Proc. NAACL (2019).

[4] J. He, D. Spokoyny, G. Neubig, and T. Berg-Kirkpatrick, Lagging inference networks and posterior collapse in variational autoencoders, Proc. ICLR (2019).

[5] I. Higgins et al., β-VAE: Learning basic visual concepts with a constrained variational framework, Proc. ICLR (2017).

[6] M. Jaderberg et al., Population based training of neural networks, arXiv:1711.09846 (2017).

[7] D. P. Kingma and M. Welling, Auto-encoding variational Bayes, Proc. ICLR (2014).

[8] D. P. Kingma et al., Improved variational inference with inverse autoregressive flow, Proc. NeurIPS (2016).

[9] Lucas, G. Tucker, R. Grosse, and M. Norouzi, Don’t blame the ELBO! A linear VAE perspective on posterior collapse, Proc. NeurIPS (2019).

[10] C. Pandarinath et al., Inferring single-trial neural population dynamics using sequential auto-encoders, Nat. Methods 15, 805 (2018).

[11] F. Pei et al., Neural Latents Benchmark ‘21, NeurIPS Datasets and Benchmarks Track (2021).

[12] B. Romera-Paredes et al., Mathematical discoveries from program search with large language models, Nature 625, 468 (2024).

[13] A. R. Sedler and A. Bhowmick, lfads-torch: A PyTorch implementation of LFADS, GitHub repository (2023).

[14] C. K. Sønderby et al., Ladder variational autoencoders, Proc. NeurIPS (2016).

